# Network pharmacology of xylazine to understand Its health consequences and develop mechanistic based remediations

**DOI:** 10.1101/2024.02.08.579475

**Authors:** Arun HS Kumar

## Abstract

**Background:** The recent raise in xylazine use disorders (XUD) in humans is a significant cause for concern as comprehensive understanding of its molecular pathology is limited and hence the ability to reverse the potential adverse effects are lacking. To address this gap, this study evaluates the dose-dependent impact of xylazine and its interactions with various potential targets, to identify an optimal reversal strategy.

**Methods:** A trichotomized (Low, medium, and high) dose, volume of distribution and predicted plasma concentration of xylazine were defined. A detailed analysis of xylazine’s network protein targets and their tissue-specific expression was performed using classical pharmacoinformatic tools. Molecular docking was used to assess the drug-target affinities and identify potential reversal agents.

**Results:** The study categorized xylazine plasma concentrations ranging from 5-8 *μ*M, 14-20 *μ*M, and 28-40 *μ*M, as low, medium, and high respectively. Xylazine displayed preferential affinity for hydrolases, kinases, transporters, and ion channels. Xylazine’s network analysis revealed the following proteins, ABCC9, RET, RAPGEF4, ACHE, TGFBR1, PGR, KCNH2, KCNN2, and TRPM8 as its high affinity targets. The tissue-specific expression of these high-affinity targets suggested potential adverse effects on various organs, particularly skeletal and smooth muscles, and the adrenal gland. The study further explored the potential reversal of xylazine pharmacology using alpha2AR-antagonists and CNS stimulants. Prazosin emerged as the most promising candidate, exhibiting a 200 to 2000-fold superior affinity against all high-affinity targets of xylazine.

**Conclusion:** This study contributes to our understanding of xylazine’s molecular mechanisms and suggests that prazosin can serve as an effective therapeutic option for mitigating xylazine-induced adverse effects in XUD patients, which warrants clinical investigation.

## Introduction

Xylazine is an alpha-2 adrenergic receptor agonist and is approved by the FDA for veterinary use as an anaesthetic.^1-3^ Unfortunately, xylazine is adulterated with illicit drugs (fentanyl, cocaine) either to amplify the effects of the drugs or boost their street value by adding weight. The prevalence of xylazine co-use with opioid/stimulant combinations is widely reported to be responsible for serious life-threatening effects (respiratory depression, hypotension, bradycardia vasoconstriction, and severe necrotic skin ulcerations/infections) leading to significant morbidity and mortality.^4-7^ Currently the mechanistic insights into the undesired effects produced by misuse of xylazine are lacking^8,9^ and therefore there aren’t any FDA-approved agents to reverse xylazine effects in humans. Additionally, the off-label use of adrenergic antagonists for emergency treatment of accidental xylazine injection have not shown acceptable clinical efficacy.^3,4,8,10^ Following extensive review of the impact of xylazine on the opioid crisis, and its growing role in overdose deaths in all regions of the United States, the White House’s Office of National Drug Control Policy has recently designated fentanyl mixed with xylazine as an emerging threat to the United States.^11,12^ Very limited information is available regarding the consequences of using drugs adulterated with xylazine and the mechanisms by which the observed undesired effects or potential substance use disorders (SUDs) are produce.^3-5,8,10^ Hence research into consequences of xylazine use which leads to undesired effects and impacts treatment of opioid abuse and overdose is necessary to develop mechanistic based optimal clinical interventions.

To address this gap in the literature, in this study network pharmacology of xylazine was evaluated to know the receptor binding profiles of xylazine and identify suitable antagonists with potential to reverse xylazine pharmacology.

## Materials and methods

The isomeric SMILES sequence of xylazine (CC1=C(C(=CC=C1)C)NC2=NCCCS2) obtained from the PubChem database was inputted into the SwissTargetPrediction server (http://www.swisstargetprediction.ch/) and STITCH database (http://stitch-db.org/) to identify the targets specific to homo sapiens.^13-15^

The pharmacokinetic parameters of xylazine were assessed using the SwissADME server and data reported in the literature. The observed dosage (mg/day), plasma concentration achieved (*μ*M) and volume of distribution (L) was trichotomized into low, medium, and high categories.

The affinity values of xylazine to all its potential targets identified were assessed using AutoDock vina 1.2.0 as reported before for other ligand-receptor combinations.^13-16^ The observed targets of xylazine were categorised based on various functional groups and the average affinity of xylazine towards each of the functional groups was estimated to assess the selectiveness of xylazine to any specific target group/s.

The high affinity targets of xylazine were identified based on a concentration affinity (CA) ratio of <0.077. The specific interaction sites, binding pocket sequence, EC_50_ values, and the number of hydrogen bonds formed during receptor-ligand interaction were also estimated as reported previously.^13-15^

The tissue specific expression pattern of the high affinity targets of xylazine were assessed using the human protein atlas (https://www.proteinatlas.org/) database. The gene code of the high affinity target was inputted into the search tool of human protein atlas database and the “tissue” tool was used to extract the protein (or mRNA) expression pattern of the target in various tissues. Only the major expression (top three tissues expressing medium/high levels of the target) was of focus in this study and are presented as Venn diagrams. The primary function of all the high affinity targets was identified from the UniProt (https://www.uniprot.org/) database.

To identify the most optimal approach to antagonise the xylazine pharmacology, in this study most know alpha-2 adrenergic receptor-antagonists (yohimbine, chlorpromazine, phentolamine, mianserine, spiperone, prazosin, alprenolol, propranolol, pindolol, atipamezole, dexmedetomidine and tolazoline) and CNS stimulants (4-aminopyridine, doxapram and caffeine) which are approved for clinical use were assessed for their affinity against all high affinity targets of xylazine as mentioned above.^13-15^ A heat map of the affinity ratio values of the agonist (xylazine) / antagonist was generated to identify the most optimal drug to antagonise the xylazine pharmacology.

## Results

The pharmacological effects of xylazine are mediated by activation of alpha-2 adrenergic receptors in the central nervous system, leading to sedation, muscle relaxation and analgesia in animals. Xylazine consists of the 2,6-dimethylphenylamino group which is attached to the fourth carbon atom of the thiazine ring (Figure 1). The nitrogen and sulphur atoms in the thiazine ring and the 2,6-dimethylphenylamino group seem to be the key components responsible for xylazine’s pharmacological properties, particularly in its interaction with alpha-2 adrenergic receptors. In this study we defined the dose of xylazine into the following three categories, 1) low (<300 mg/day, observed range 150 to 300 mg/day), 2) medium (>300 and <800 mg/day) and 3) high (>800 mg/day, observed range 800 to 1200 mg/day) (Figure 1). These dose ranges were used to estimate the plasma concentrations achievable in humans, which ranged from 5-8 *μ*M (low dose), 14-20 *μ*M (medium dose) and 28-40 *μ*M (high dose) by considering the following variable volume of distribution of xylazine in humans [110 to 120 litres (low), 135 to 145 litres (medium) and 155 to 165 litres (high)] (Figure 1).

**Figure 1:**
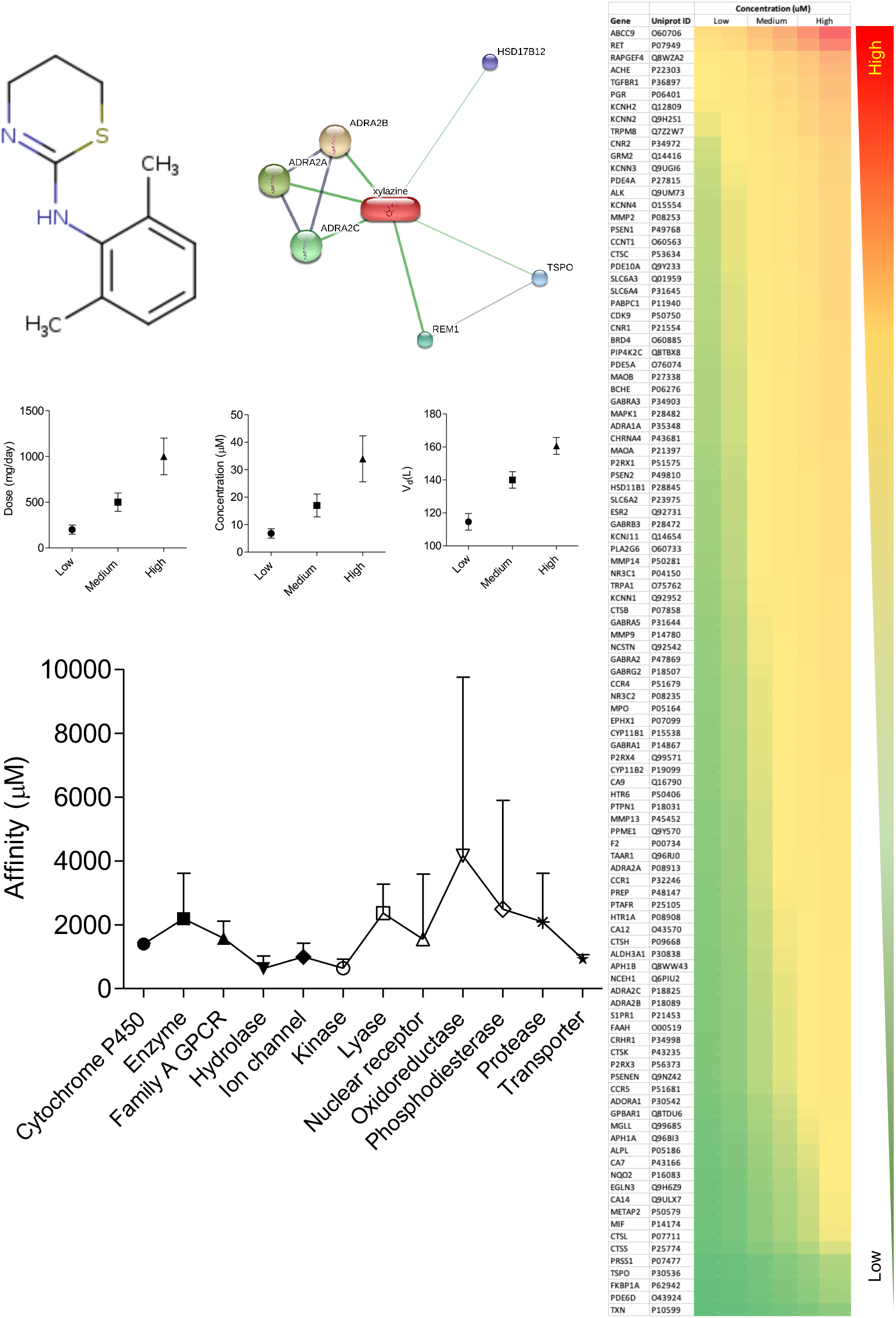
Pharmacological properties of xylazine. Chemical structure of xylazine along with its network proteins identified in the STITCH database. The middle graphs show the trichotomized (low, medium, and high) data of xylazine dose (mg/day), predicted plasma concentration (*μ*M) and volume of distribution (L) in humans. The heat map represents the concentration affinity ratio of xylazine against all its identified targets from the SwissTargetPrediction database (Scale: red to green = high to low ratio values). The bottom graph represents affinity (*μ*M, mean ± SD) of xylazine against all its target categories identified from the SwissTargetPrediction database.

The xylazine network analysis in the STITCH database showed six network proteins, of which four were close networks (affinity scores >0.8) consisting of alpha-2 adrenergic receptors (types 2A, 2B and 2C) and RAS (RAD and GEM)-like GTP-binding 1 (REM1), while the other two proteins showed weaker network (affinity scores <0.5), consisting of hydroxysteroid (17-beta) dehydrogenase 12 (HSD17B12) and translocator protein (TSPO) (Figure 1). The network analysis of xylazine in the SwissTargetPrediction server showed 105 potential targets. The affinity of the xylazine to these targets ranged from 147 to 10616 *μ*M). The following nine targets, ABCC9, RET, RAPGEF4, ACHE, TGFBR1, PGR, KCNH2, KCNN2 and TRPM8 had affinity values of <500 *μ*M with xylazine and were also recognised has high affinity targets based on their concentration affinity (CA) ratio of <0.077 (Figure 1). The highest affinity of xylazine was observed for ABCC9 (147 *μ*M). To assess if xylazine has preferential affinity to any specific target class, the 105 potential targets were grouped into 12 target class and the average affinity of xylazine to each of the target class was estimated and is summarised in figure 1. The highest affinity of xylazine was observed for hydrolases, kinases, transporters, and ion channels, this was followed by affinity for Family A GPCR and cytochrome P_450_ (Figure 1). The least affinity of xylazine was observed for oxidoreductases (Figure 1).

Incidentally the alpha-2 adrenergic receptors (ADRA2A, ADRA2B and ADRA2C) were not among the high affinity targets of xylazine with affinity values ranging from 1590 to 1921 *μ*M. Hence EC_50_ values (extrapolated from the binding energies) of xylazine against all its high affinity targets and the alpha-2 adrenergic receptors was evaluated (Figure 2). The least EC_50_ of xylazine was observed for PGR, followed by TRPM8 and ADRA2A (Figure 2). The specific binding sequence of xylazine targets identified in this study together with the number of hydrogen bonds formed during the interaction are summarised in figure 2. The correlation between the EC_50_ values and the number of hydrogen bonds formed between xylazine, and its high affinity targets suggested a very weak correlation (r^2^<0.016) (data not shown). The weak correlation between binding affinity and hydrogen bonding pattern is consistent with the intricate nature of ligand-receptor interactions. The highest number of hydrogen bonds (>18) between xylazine and its high affinity targets was observed for TGFBR1, ADRA2A, ABCC9, ADRA2B, and ACHE (Figure 2). While the number of hydrogen bonds formed between xylazine and RAPGEF4, PGR, TRPM8, and ADRA2C ranged from 13-15 bonds. RET, KCNH2, and KCNN2 showed the least number (<10) of hydrogen bonds formed with xylazine.

**Figure 2:**
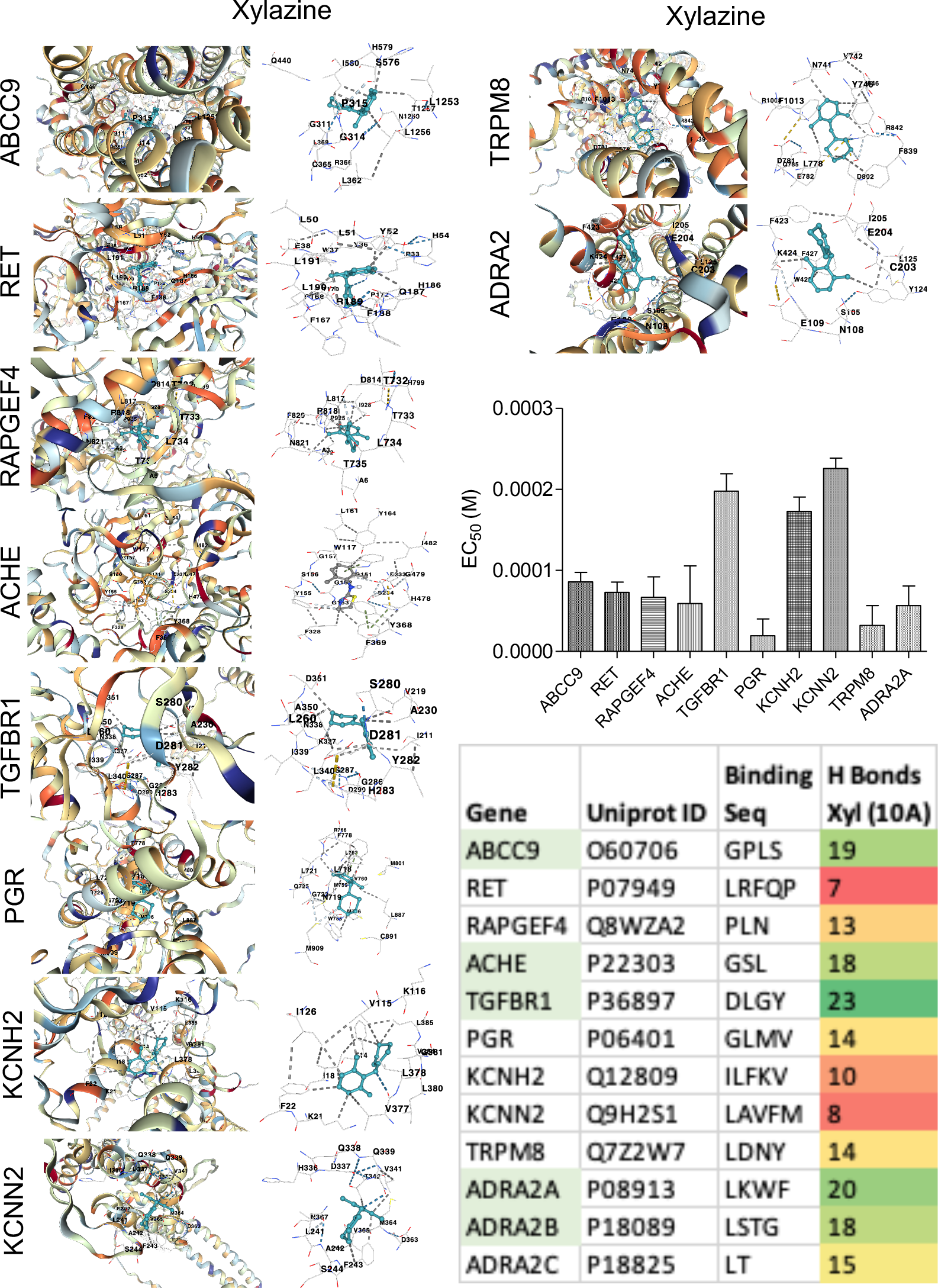
Xylazine and its high affinity targets. Representative images from molecular docking showing interaction of xylazine with its high affinity targets. The bar graph represents the EC_50_ values (molar (M), mean ± SD) of xylazine for its high affinity targets. Inset table shows the amino acid sequence of the binding pockets of the xylazine’s high affinity targets along with the number of hydrogen bonds formed at 10Å distance (the targets forming ≥ 18 hydrogen bonds are highlighted in light green).

The tissue specific expression of all the high affinity targets of xylazine were assessed from the human protein atlas database and are summarised in figure 3. The targets of xylazine which formed the most number of hydrogen bonds (>18) were mainly expressed in bone marrow, cerebellum, colangiocytes, heart, hypothalamus, ionocytes, langerhans cells, liver, paneth cells, skeletal muscle, and smooth muscle (Figure 3). Of these xylazine targets, ABCC9, ACHE, TGFBR1 and ADRA2C were predominantly expressed on skeletal and smooth muscles (Figure 3). While RET, RAPGEF4, KCNN2 were considerably expressed on adrenal gland (Figure 3), suggesting that the major adverse effects of xylazine could be consequence to compromised physiology of adrenal gland and skeletal and smooth muscles. To identify the drug which can optimally antagonise xylazine pharmacology, all its high affinity targets were screened for affinity against selected alpha2AR-antagonists and CNS stimulants which are clinically used (Figure 3).

**Figure 3:**
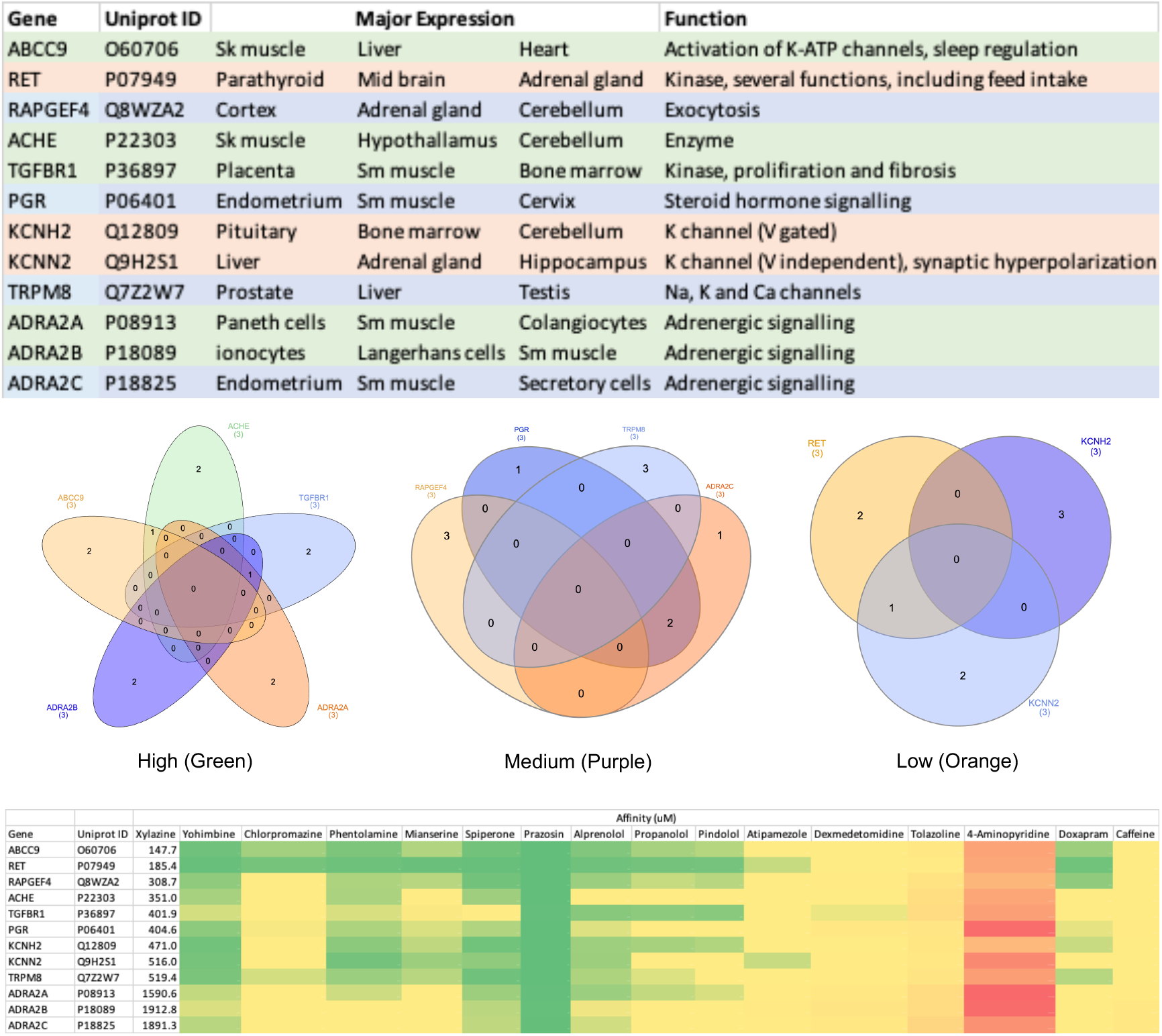
Expression and antagonism of high affinity targets of xylazine. The major expression of (top three tissues) high affinity targets of xylazine are shown along with their major function. The Venn diagrams trichotomize (High: Green, medium: purple, and low: orange) the expression pattern of xylazine targets based on their affinities. The heat map represents the affinity ratios of xylazine and its antagonist to the high affinity targets of xylazine (scale: green represents lowest ratio indicating better antagonism, while red represents highest ratio indicating poor antagonism).

A ratio of the affinities of xylazine and selected antagonist to the specific target was estimated to assess the relative potential of the drug to reverse xylazine pharmacology (Figure 3). In this screening, prazosin was identified as the most optimal drug to reverse xylazine pharmacology with ∼ 200 to 2000 (average 660) fold superior affinity against all high affinity targets of xylazine. This was followed by spiperone (26-fold), yohimbine (21-fold), phentolamine (14-fold) and alprenolol (13-fold) (Figure 3).

## Discussion

The results of this study contribute significantly to the understanding of xylazine’s pharmacological profile and has provides insights into optimally reversing its effects in humans. The comprehensive investigation into the pharmacological profile of xylazine has unveiled details of its dose-dependent pharmacokinetics, molecular interactions, potential adverse effects, and identified promising reversal agent/s. This study expands beyond the pharmacological effects of xylazine mediated by activation of alpha-2 adrenergic receptors and offers knowhow on possibilities of xylazine interacting with other targets with much superior affinity in humans. There are two major reasons to justify the study observations, 1) In human patients with xylazine use disorders (XUDs), the clinical symptoms observed are not classical of alpha-2 adrenergic receptor activation^4,6^ and 2) The clinical approach to correcting adverse effects in XUD patients using alpha-2 adrenergic receptor antagonist show very poor efficacy.^1,4,9^ These two reasons justify that the adverse effects observed in XUDs are mediated possibly by receptors other than alpha-2 adrenergic receptors and this study specifically identifies nine targets (ABCC9, RET, RAPGEF4, ACHE, TGFBR1, PGR, KCNH2, KCNN2 and TRPM8) which can be involved in XUDs.

These specific targets of XUD identified offer a platform to develop effective and targeted reversal agents.

The delineation of xylazine doses into low, medium, and high categories, along with the associated estimation of plasma concentrations in humans, has significant implications for comprehending reversal of xylazine pharmacology. Recognizing how different doses impact plasma concentrations is fundamental for devising effective reversal protocols, specifically in the context of concentration affinity ratios of the drugs. The plasma concentration ranges defined in this study are consistent with previous reports^1,4,9^ and offers valuable insights into the pharmacokinetics/dynamics of xylazine and its reversal in the human body. This information becomes pivotal for clinicians seeking to design protocols that efficiently and safely counteract the adverse effects of xylazine. Categorizing xylazine doses also allows for an assessment of the potential risks associated with reversal at higher doses, where plasma concentrations are elevated to influence many more targets. This understanding is crucial for balancing the efficacy of reversal agents with safety considerations for the patient.^3,17,18^ This knowledge also aids in optimizing the choice and dosage of reversal agents to achieve prompt and effective reversal while minimizing adverse effects. This study by characterization of xylazine doses and their corresponding plasma concentrations, considering variable volumes of distribution, significantly enhances the understanding of reversing xylazine pharmacology in humans. This information serves as a cornerstone for developing targeted and safe reversal protocols, ensuring effective mitigation of xylazine-induced adverse effects while prioritizing patient well-being. The observed ranges in this study also offer a foundation for future research on the pharmacokinetics of xylazine in diverse patient populations.

The potential targets of xylazine identified in this study highlight its pharmacological profile, which is crucial to assessing the major adverse effects of xylazine reported in human XUD patients.^6,8,10,19^ While the pharmacological effects of xylazine through the adrenergic receptors are well known, this study identifies several significantly higher affinity targets of xylazine, which are previously not reported. The highest average-affinities of xylazine to target categories such as hydrolases, kinases, transporters, and ion channels suggest a broad impact on several cellular processes potentially regulated by ABCC9, RET, RAPGEF4, ACHE, TGFBR1, PGR, KCNH2, KCNN2, and TRPM8. Of these, the least EC_50_ of xylazine was observed for PGR, followed by TRPM8, ADRA2A, ABCC9, RET, RAPGEF4 and ACHE. The interplay between xylazine and alpha-2 adrenergic receptors, reinforces the established role of these receptors in mediating sedation and analgesia.^1,2,20,21^ While the involvement of TRPM8, ADRA2A, ABCC9, RET, RAPGEF4 and ACHE as xylazine targets expands our understanding of potential signalling cascades beyond direct adrenergic receptor interactions.^1,2,17,20^ This interconnected network underscores the need for a holistic approach when studying the pharmacodynamics of xylazine. Identification of high-affinity targets such as TRPM8, ADRA2A, ABCC9, RET, RAPGEF4 and ACHE aligns with recent reports on adverse effects associated with xylazine abuse, which are not classical of alpha-2 adrenergic receptor stimulation.^5,6,18,19,21^ The tissue-specific expression of these targets, especially in the liver, adrenal gland and skeletal/smooth muscles, is also consistent with clinical observations of xylazine’s impact on microvasculature, endocrine and musculoskeletal systems.^4-6,8,18,19,21^ This correlation supports the notion that the undesired effects of xylazine can be attributed to its interactions with the specific targets identified in this study. The observation of xylazine interaction with KCNN2 and KCNH2 are consistent with most clinical reports of adverse cardiovascular effects in XUD patients.^4,9^ Xylazine was also observed to target ACHE, which can lead to enhanced acetylcholine levels, this together with its sympatholytic effects^1,3^ can severely imbalance autonomic physiology leading to compromised cardiovascular and respiratory functions, which are consistently observed in XUD patients.^1,3,4,8^ XUD patients also suffer from altered sleep patterns^1,3,4^ and in this study the identification of ABCC9 as a target of xylazine provides a mechanistic insight into this pharmacological effects observed. ABCC9, which is a subunit of ATP-sensitive potassium channels is known to regulate sleep duration^22-25^ possibly by influencing the diffuse modulatory system^25,26^ through compromised functions of RET and RAPGEF4,^27-29^ which are also high affinity targets of xylazine. In addition to the effects of xylazine on autonomic, endocrine and musculoskeletal systems, it is likely to influence sensory physiology as well by targeting TRPM8, which is a receptor-activated non-selective cation channel involved in detection of cold (< 25°C) and intracellular pH.^30,31^ Such broad spectrum pharmacology of xylazine pose several challenges when treating its adverse effects. The involvement of multiple targets of xylazine in its observed adverse effects, necessitate optimally targeting them for effectively reversing its effects. Identification of such an approach is only possible through network pharmacology analysis.^13-15^ The results of the screening for high-affinity targets of xylazine and their antagonism by selected alpha-2 adrenergic receptor-antagonists and CNS stimulants provide valuable insights into potential candidates for reversing xylazine pharmacology. In this study prazosin emerged as the most optimal drug for reversing xylazine pharmacology, demonstrating a remarkable 200 to 2000-fold superior affinity against all high-affinity targets of xylazine. Hence prazosin could be a highly effective antagonist for mitigating the effects of xylazine, making it a promising candidate for further investigation and potential clinical use. The identification of prazosin as a potent antagonist aligns with previous research on alpha-2 adrenergic receptor antagonists, emphasizing their potential clinical applications^22,32^ and this study further expands on the clinical utility of prazosin through its effects independent of alpha-2 adrenergic receptors. The currently used treatment options for xylazine toxicity include, various combinations of intravenous (IV) fluids, saline ocular irrigation, mechanical ventilation, saline cathartics, gastric lavage, silver sulfadiazine cream / antibacterial ointments and/or yohimbine / tolazoline / atipamezole / naloxone (1.2 to 2 mg), lidocaine / atropine / magnesium infusion / activated charcoal / metoprolol succinate / thiamine / clonidine, all of which have shown only limited clinical efficacy in XUD patients.^1,3,4,6,9^ Considering the current clinical limitations, the potential use of prazosin in reversing xylazine pharmacology suggested in this study offers a detailed mechanistic insights which merits its clinical evaluation. In comparison with other antagonists evaluated in this study, prazosin stands out as a particularly potent antagonist and hence evaluating its efficacy in XUD patients should be considered to validate prazosin’s effectiveness in real-world scenarios and explore its safety profile in diverse patient populations.

In summary, this study provides a robust mechanistic understanding of the pharmacological properties of xylazine, offering insights into potential adverse effects and avenues for targeted interventions. The identified high-affinity targets, network interactions, and the promising role of prazosin as a reversal agent set the stage for future investigations in XUD patients.

## Acknowledgements

Research support from University College Dublin-Seed funding/Output Based Research Support Scheme (R19862, 2019), Royal Society-UK (IES\R2\181067, 2018) and Stemcology (STGY2917, 2022) is acknowledged.

